# A general evolution landscape of language and cognition genes

**DOI:** 10.1101/2024.03.11.584338

**Authors:** Zhizhou Zhang, Shuaiyu Zhang, Hongjie Zhou, Yongdong Xu

## Abstract

The polymorphism profiles of language genes (LG) display different patterns across various ancient and modern populations, leading to the speculation that cognition gene (CG) polymorphism profiles may exhibit similar trends. However, the evolutionary processes of language gene polymorphism patterns (LGPP) and cognition gene polymorphism patterns (CGPP) are likely to demonstrate distinct characteristics. In particular, it is intriguing to determine whether there is any overlap in the timing of significant changes in CGPP and LGPP over the large timescales of evolution. The potential existence of such overlap can also be assessed by examining whether the samples carrying significant changes in LGPP and CGPP are the same. This study investigated the genetic differences at 239 SNP loci in 18 language genes (LG) and 223 SNP loci in 18 cognition genes (CG) across 170 whole genomes. Principal component analysis (PCA) was used to cluster the SNP data of the aforementioned samples, and the similarity of SNP patterns between each sample was calculated from three perspectives: LG, CG, and CGLG. The basic conclusions are as follows: (1) If different positions in the PCA analysis results can essentially represent the pattern differences in SNP polymorphisms, then both language gene polymorphism patterns and cognition gene polymorphism patterns have undergone distinct stages of evolution; (2) There were significant differences in the early manifestations of language gene polymorphism patterns and cognition gene polymorphism patterns during human evolution: Language gene polymorphism patterns could not differentiate general animals, primates, and ancient human samples in the early stages of evolution, whereas cognition gene polymorphism patterns seemed to be initially divisible into two patterns, one closely resembling a group of animals and certain ancient human samples, and the other reflected in a different set of animal and primate samples. (3) It appears that samples from all five continents can be observed at every stage of evolution, suggesting that new evolving populations have always had ample time to spread across continents. (4) A quantitative comparison of the SNP profiles of 170 samples revealed that their CG and LG plus CGLG profiles indeed have 2-3 potential significant change points, and the samples carrying these significant change points has 2 common samples, namely ge1 (Georgia) and us2 (North America), implying that the most significant changes in language or cognition gene polymorphism patterns during human evolution may have occurred in some human populations in Europe/ North America.

## Indroduction

There are various, yet complementary, theories about the evolution of human language [1-6]. Clearly, all animals have their own ways of communication, even if not always through vocal sounds made with the mouth. Primates, for example, have at least a dozen distinct vocal sounds that carry specific meanings, which can be considered a rudimentary form of language. Some studies strongly believe in the following hypothesis: once the primate brain evolved to a certain stage and suddenly acquired symbolic thinking ability [5-6], Homo sapiens came into being. Symbolic thinking naturally possesses the capacity to gradually refine and complicate the meanings of language. And once language could become more complex and precise in meaning, and its significance could be passed down, humans were able to progressively accumulate their ancestors’ experiences, accelerating human evolution.

The above process also implies several facts: 1) The evolution of language occurred much earlier than the ability for symbolic thinking. The capacity for language is primarily a capability of vocal sounds, resulting from the movement of muscles inside the body such as those in the mouth, and is fundamentally a type of motor skill; (2) Symbolic thinking is not a uniquely human cognitive ability, but there are varying degrees of symbolic thinking. Ordinary animals, even if they appear relatively intelligent, cannot compare with human symbolic thinking. If we disregard the levels of symbolic thinking and simply measure its presence or absence, then this point in time likely occurred about 35,000∼70,000 years ago [5,7-8]. If we use different levels to measure symbolic thinking, we can better understand why human evolution is divided into stages such as ancient apes, hominids, Homo erectus, Homo habilis, Homo sapiens, and modern humans; (3) Language and cognitive abilities complement each other, so the period following the emergence of Homo sapiens in the evolutionary process (which could span tens of thousands to hundreds of thousands of years) should have been a time of rapid advancement in both language and cognitive abilities. Through the proliferation, interbreeding, and iteration of populations, most human groups on Earth gradually came to possess advanced language and cognitive abilities, while those groups and individuals with only one of these advanced abilities would accelerate towards extinction. (4) Due to the vast and diverse geographical environments on Earth, various conditions are provided for the long-term existence of specific populations, so there is still, in principle, genetic diversity in language and cognitive abilities on Earth. Even in large cities, or to say, around us, we can still encounter individuals with severe language impairments but outstanding cognitive abilities. There are also many individuals who are proficient in spoken language but have particularly weak symbolic thinking abilities (such as in mathematics). Some children nearing the age of 10 still do not possess adequate language skills. These are probably all specific intermediate states in the evolutionary process of the two abilities mentioned above. (5) The nature of language determines that it only has clear meanings within specific contexts, which aligns with the current situation where there are over 7,500 dialects worldwide. However, human evolution has far surpassed these limitations. Humans can now design entirely new languages independent of any local context and teach and spread them anywhere. (6) As the brain’s spatial structure increasingly shows clear correspondence and working principles for language and cognitive abilities, detailed quantitative measurement of these abilities in principle allows for definitive diagnosis of brain diseases, aging, etc., and forms the basis for developing new brain-computer interface technologies and products.

The process of language evolution and cognitive evolution should be reflected in the human genome sequence, and the modern human genome should also contain some of this evolutionary information. The older the fossil DNA, the more likely it is to reveal a greater number of intermediate states of evolution. Moreover, in principle, one should be able to observe different evolutionary rhythms for language genes and cognition genes along the evolutionary path. This study attempts to analyze a set of linguistic gene polymorphisms and cognitive gene polymorphisms in a batch of ancient DNA and modern genome sequences, thereby depicting a general landscape of the evolution of language and cognition genes.

## Methods

### Genome sequences

Genome sequences were downloaded from ENA database (https://www.ebi.ac.uk/ena/browser/), SRA database (https://www.ncbi.nlm.nih.gov/sra) and Ensembl genome browser. Total 170 whole genomes (including 59 ancient genomes, Table 1) from 5 continents (Africa, Asia, Europe, North America, and South America) were collected. The above genome sequences have fastq, fn or fna formats, and all can be read and scanned with python based hash07plus03 software.

**Table 1.**
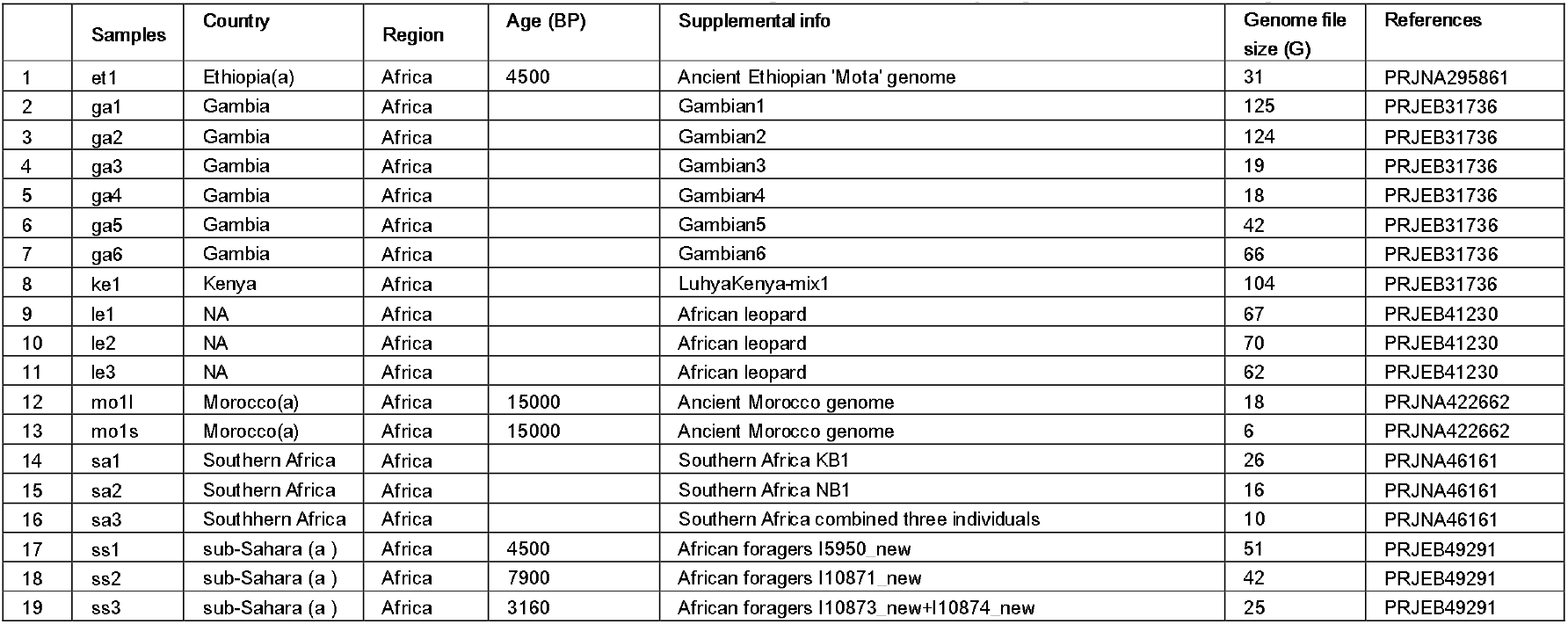
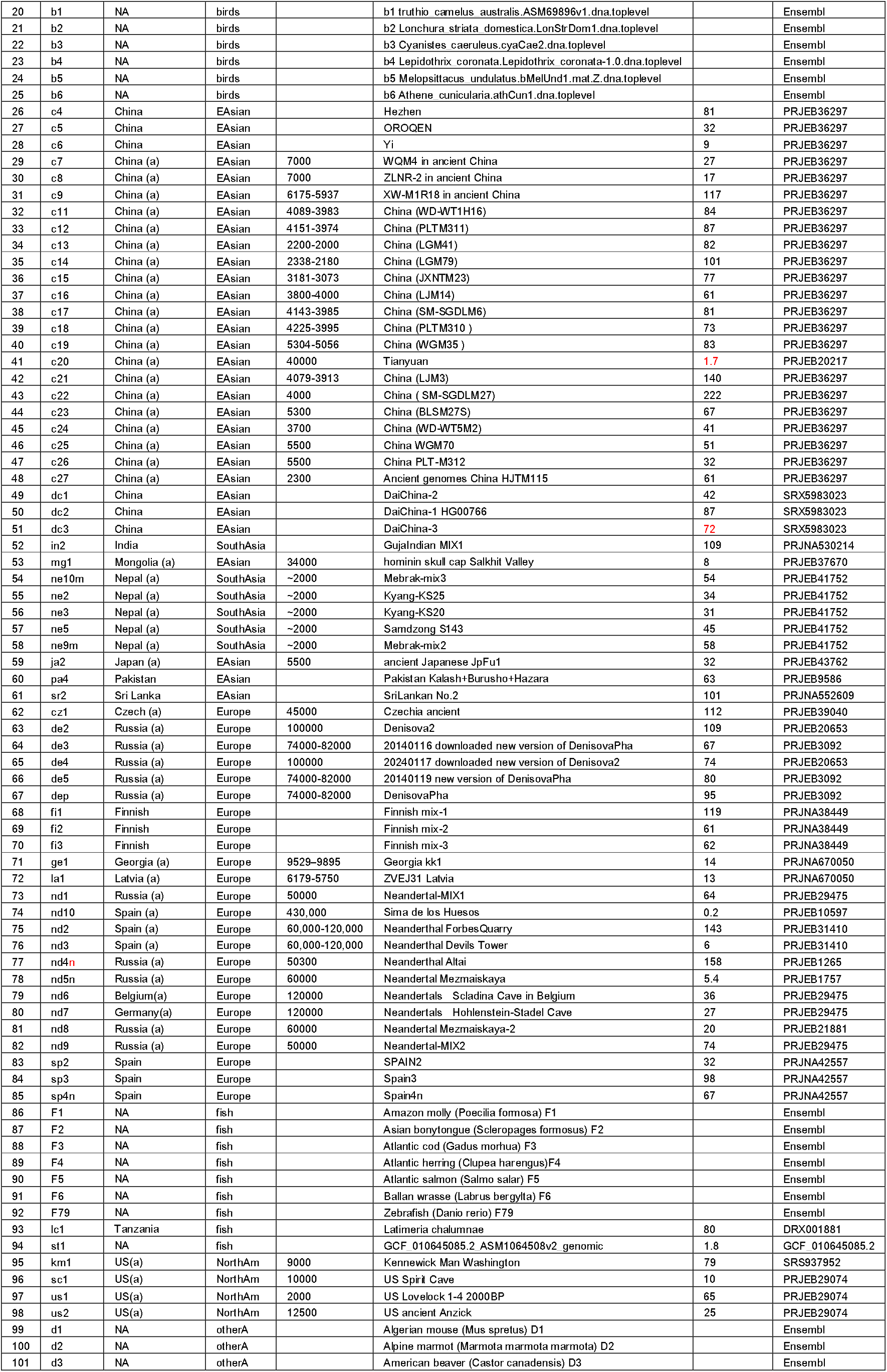
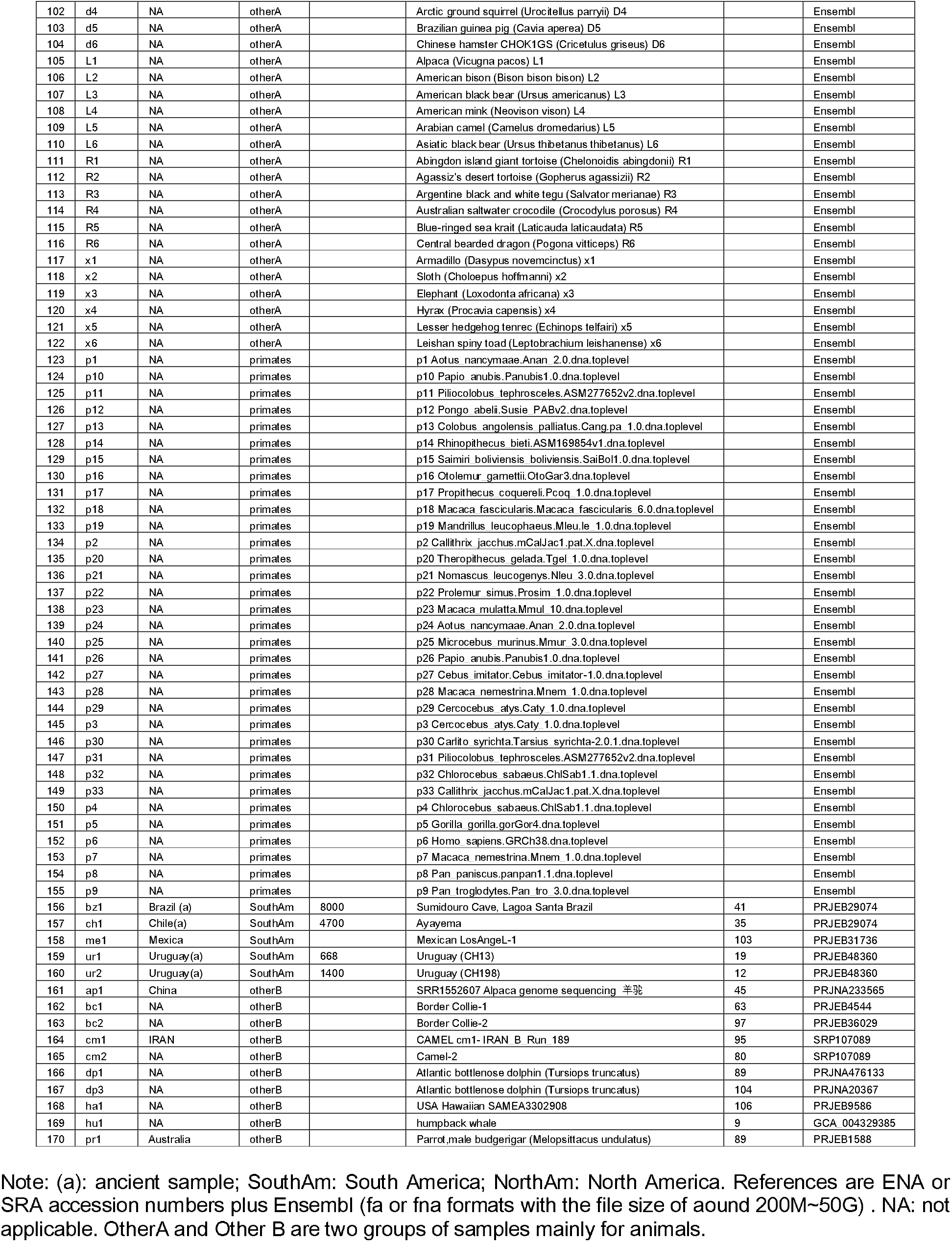
The 170 whole genomes employed in this study.

|  | Samples | Country | Region | Age (BP) | Supplemental info | Genome file size (G) | References |
| --- | --- | --- | --- | --- | --- | --- | --- |
| 1 | et1 | Ethiopia (a) | Africa | 4500 | Ancient Ethiopian 'Mota' genome | 31 | PRJNA295861 |
| 2 | ga1 | Gambia | Africa |  | Gambian1 | 125 | PRJEB31736 |
| 3 | ga2 | Gambia | Africa |  | Gambian2 | 124 | PRJEB31736 |
| 4 | ga3 | Gambia | Africa |  | Gambian3 | 19 | PRJEB31736 |
| 5 | ga4 | Gambia | Africa |  | Gambian4 | 18 | PRJEB31736 |
| 6 | ga5 | Gambia | Africa |  | Gambian5 | 42 | PRJEB31736 |
| 7 | ga6 | Gambia | Africa |  | Gambian6 | 66 | PRJEB31736 |
| 8 | ke1 | Kenya | Africa |  | LuhyaKenya-mix1 | 104 | PRJEB31736 |
| 9 | le1 | NA | Africa |  | African leopard | 67 | PRJEB41230 |
| 10 | le2 | NA | Africa |  | African leopard | 70 | PRJEB41230 |
| 11 | le3 | NA | Africa |  | African leopard | 62 | PRJEB41230 |
| 12 | mo1l | Morocco (a) | Africa | 15000 | Ancient Morocco genome | 18 | PRJNA422662 |
| 13 | mo1s | Morocco (a) | Africa | 15000 | Ancient Morocco genome | 6 | PRJNA422662 |
| 14 | sa1 | Southern Africa | Africa |  | Southern Africa KB1 | 26 | PRJNA46161 |
| 15 | sa2 | Southern Africa | Africa |  | Southern Africa NB1 | 16 | PRJNA46161 |
| 16 | sa3 | Southern Africa | Africa |  | Southern Africa combined three individuals | 10 | PRJNA46161 |
| 17 | ss1 | sub-Saharan (a) | Africa | 4500 | African foragers I5950_new | 51 | PRJEB49291 |
| 18 | ss2 | sub-Saharan (a) | Africa | 7900 | African foragers I10871_new | 42 | PRJEB49291 |
| 19 | ss3 | sub-Saharan (a) | Africa | 3160 | African foragers I10873_new+I10874_new | 25 | PRJEB49291 |
| 20 | b1 | NA | birds |  | b1 truthio_camelus_australis.ASM69896v1.dna.toplevel |  | Ensembl |
| 21 | b2 | NA | birds |  | b2 Lonchura_striata_domestica.LonStrDom1.dna.toplevel |  | Ensembl |
| 22 | b3 | NA | birds |  | b3 Cyanistes_caeuleus.cyaCae2.dna.toplevel |  | Ensembl |
| 23 | b4 | NA | birds |  | b4 Lepidothrix_coronata.Lepidothrix_coronata-1.0.dna.toplevel |  | Ensembl |
| 24 | b5 | NA | birds |  | b5 Melopsittacus_undulatus.bMelUnd1.mat.Z.dna.toplevel |  | Ensembl |
| 25 | b6 | NA | birds |  | b6 Athene_unicularia.athCun1.dna.toplevel |  | Ensembl |
| 26 | c4 | China | EAsian |  | Hezhen | 81 | PRJEB36297 |
| 27 | c5 | China | EAsian |  | OROQEN | 32 | PRJEB36297 |
| 28 | c6 | China | EAsian |  | Yi | 9 | PRJEB36297 |
| 29 | c7 | China (a) | EAsian | 7000 | WQM4 in ancient China | 27 | PRJEB36297 |
| 30 | c8 | China (a) | EAsian | 7000 | ZLNR-2 in ancient China | 17 | PRJEB36297 |
| 31 | c9 | China (a) | EAsian | 6175-5937 | XW-M1R18 in ancient China | 117 | PRJEB36297 |
| 32 | c11 | China (a) | EAsian | 4089-3983 | China (WD-WT1H16) | 84 | PRJEB36297 |
| 33 | c12 | China (a) | EAsian | 4151-3974 | China (PLTM311) | 87 | PRJEB36297 |
| 34 | c13 | China (a) | EAsian | 2200-2000 | China (LGM41) | 82 | PRJEB36297 |
| 35 | c14 | China (a) | EAsian | 2338-2180 | China (LGM79) | 101 | PRJEB36297 |
| 36 | c15 | China (a) | EAsian | 3181-3073 | China (JXNTM23) | 77 | PRJEB36297 |
| 37 | c16 | China (a) | EAsian | 3800-4000 | China (LJM14) | 61 | PRJEB36297 |
| 38 | c17 | China (a) | EAsian | 4143-3985 | China (SM-SGDLM6) | 81 | PRJEB36297 |
| 39 | c18 | China (a) | EAsian | 4225-3995 | China (PLTM310) | 73 | PRJEB36297 |
| 40 | c19 | China (a) | EAsian | 5304-5056 | China (WGM35) | 83 | PRJEB36297 |
| 41 | c20 | China (a) | EAsian | 40000 | Tianyuan | 1.7 | PRJEB20217 |
| 42 | c21 | China (a) | EAsian | 4079-3913 | China (LJM3) | 140 | PRJEB36297 |
| 43 | c22 | China (a) | EAsian | 4000 | China (SM-SGDLM27) | 222 | PRJEB36297 |
| 44 | c23 | China (a) | EAsian | 5300 | China (BLSM27S) | 67 | PRJEB36297 |
| 45 | c24 | China (a) | EAsian | 3700 | China (WD-WT5M2) | 41 | PRJEB36297 |
| 46 | c25 | China (a) | EAsian | 5500 | China WGM70 | 51 | PRJEB36297 |
| 47 | c26 | China (a) | EAsian | 5500 | China PLT-M312 | 32 | PRJEB36297 |
| 48 | c27 | China (a) | EAsian | 2300 | Ancient genomes China HJTM115 | 61 | PRJEB36297 |
| 49 | dc1 | China | EAsian |  | DaiChina-2 | 42 | SRX5983023 |
| 50 | dc2 | China | EAsian |  | DaiChina-1 HG00766 | 87 | SRX5983023 |
| 51 | dc3 | China | EAsian |  | DaiChina-3 | 72 | SRX5983023 |
| 52 | in2 | India | SouthAsia |  | GujarIndian MIX1 | 109 | PRJNA530214 |
| 53 | mg1 | Mongolia (a) | EAsian | 34000 | hominin skull cap Salkhit Valley | 8 | PRJEB37670 |
| 54 | ne10m | Nepal (a) | SouthAsia | ~2000 | Mebrak-mix3 | 54 | PRJEB41752 |
| 55 | ne2 | Nepal (a) | SouthAsia | ~2000 | Kyang-KS25 | 34 | PRJEB41752 |
| 56 | ne3 | Nepal (a) | SouthAsia | ~2000 | Kyang-KS20 | 31 | PRJEB41752 |
| 57 | ne5 | Nepal (a) | SouthAsia | ~2000 | Samdzong S143 | 45 | PRJEB41752 |
| 58 | ne9m | Nepal (a) | SouthAsia | ~2000 | Mebrak-mix2 | 58 | PRJEB41752 |
| 59 | ja2 | Japan (a) | EAsian | 5500 | ancient Japanese JpFu1 | 32 | PRJEB43762 |
| 60 | pa4 | Pakistan | EAsian |  | Pakistan Kalash+Burusho+Hazara | 63 | PRJEB9586 |
| 61 | sr2 | Sri Lanka | EAsian |  | SriLankan No 2 | 101 | PRJNA552609 |
| 62 | cz1 | Czech (a) | Europe | 45000 | Czechia ancient | 112 | PRJEB39040 |
| 63 | de2 | Russia (a) | Europe | 100000 | Denisova2 | 109 | PRJEB20653 |
| 64 | de3 | Russia (a) | Europe | 74000-82000 | 20140116 downloaded new version of DenisovaPha | 67 | PRJEB3092 |
| 65 | de4 | Russia (a) | Europe | 100000 | 20240117 downloaded new version of Denisova2 | 74 | PRJEB20653 |
| 66 | de5 | Russia (a) | Europe | 74000-82000 | 20140119 new version of DenisovaPha | 80 | PRJEB3092 |
| 67 | dep | Russia (a) | Europe | 74000-82000 | DenisovaPha | 95 | PRJEB3092 |
| 68 | fi1 | Finnish | Europe |  | Finnish mix-1 | 119 | PRJNA38449 |
| 69 | fi2 | Finnish | Europe |  | Finnish mix-2 | 61 | PRJNA38449 |
| 70 | fi3 | Finnish | Europe |  | Finnish mix-3 | 62 | PRJNA38449 |
| 71 | ge1 | Georgia (a) | Europe | 9529-9895 | Georgia kk1 | 14 | PRJNA670050 |
| 72 | la1 | Latvia (a) | Europe | 6179-5750 | ZVEJ31 Latvia | 13 | PRJNA670050 |
| 73 | nd1 | Russia (a) | Europe | 50000 | Neandertal-MIX1 | 64 | PRJEB29475 |
| 74 | nd10 | Spain (a) | Europe | 430,000 | Sima de los Huesos | 0.2 | PRJEB10597 |
| 75 | nd2 | Spain (a) | Europe | 60,000-120,000 | Neandertal Forbes Quarry | 143 | PRJEB31410 |
| 76 | nd3 | Spain (a) | Europe | 60,000-120,000 | Neandertal Devils Tower | 6 | PRJEB31410 |
| 77 | nd4n | Russia (a) | Europe | 50300 | Neandertal Altai | 158 | PRJEB1265 |
| 78 | nd5n | Russia (a) | Europe | 60000 | Neandertal Mezmaiskaya | 5.4 | PRJEB1757 |
| 79 | nd6 | Belgium(a) | Europe | 120000 | Neandertals Sceldina Cave in Belgium | 36 | PRJEB29475 |
| 80 | nd7 | Germany(a) | Europe | 120000 | Neandertals Hohlenstein-Stadel Cave | 27 | PRJEB29475 |
| 81 | nd8 | Russia (a) | Europe | 60000 | Neandertal Mezmaiskaya-2 | 20 | PRJEB21881 |
| 82 | nd9 | Russia (a) | Europe | 50000 | Neandertal-MIX2 | 74 | PRJEB29475 |
| 83 | sp2 | Spain | Europe |  | SPAIN2 | 32 | PRJNA42557 |
| 84 | sp3 | Spain | Europe |  | Spain3 | 98 | PRJNA42557 |
| 85 | sp4n | Spain | Europe |  | Spain4n | 67 | PRJNA42557 |
| 86 | F1 | NA | fish |  | Amazon molly (Poecilia formosa) F1 |  | Ensembl |
| 87 | F2 | NA | fish |  | Asian bonytongue (Scleropages formosus) F2 |  | Ensembl |
| 88 | F3 | NA | fish |  | Atlantic cod (Gadus morhua) F3 |  | Ensembl |
| 89 | F4 | NA | fish |  | Atlantic herring (Clupea harengus) F4 |  | Ensembl |
| 90 | F5 | NA | fish |  | Atlantic salmon (Salmo salar) F5 |  | Ensembl |
| 91 | F6 | NA | fish |  | Ballan wrasse (Labrus bergylta) F6 |  | Ensembl |
| 92 | F79 | NA | fish |  | Zebrafish (Danio rerio) F79 |  | Ensembl |
| 93 | lc1 | Tanzania | fish |  | Latimeria chalumnae | 80 | DRX001881 |
| 94 | sl1 | NA | fish |  | GCF_010645085.2_ASM1064508v2_genomic | 1.8 | GCF_010645085.2 |
| 95 | km1 | US(a) | NorthAm | 9000 | Kennewick Man Washington | 79 | SRS937952 |
| 96 | sc1 | US(a) | NorthAm | 10000 | US Spirit Cave | 10 | PRJEB29074 |
| 97 | us1 | US(a) | NorthAm | 2000 | US Lovelock 1-4 2000BP | 65 | PRJEB29074 |
| 98 | us2 | US(a) | NorthAm | 12500 | US ancient Anzick | 25 | PRJEB29074 |
| 99 | d1 | NA | otherA |  | Algerian mouse (Mus spretus) D1 |  | Ensembl |
| 100 | d2 | NA | otherA |  | Alpine marmot (Marmota marmota marmota) D2 |  | Ensembl |
| 101 | d3 | NA | otherA |  | American beaver (Castor canadensis) D3 |  | Ensembl |
| 102 | d4 | NA | otherA |  | Arctic ground squirrel (*Urocitellus parryi*) D4 |  | Ensembl |
| 103 | d5 | NA | otherA |  | Brazilian guinea pig (*Cavia aperea*) D5 |  | Ensembl |
| 104 | d6 | NA | otherA |  | Chinese hamster CHOK1GS (*Cricetulus griseus*) D6 |  | Ensembl |
| 105 | L1 | NA | otherA |  | Alpaca (*Vicugna pacos*) L1 |  | Ensembl |
| 106 | L2 | NA | otherA |  | American bison (*Bison bison*) L2 |  | Ensembl |
| 107 | L3 | NA | otherA |  | American black bear (*Ursus americanus*) L3 |  | Ensembl |
| 108 | L4 | NA | otherA |  | American mink (*Neovison vison*) L4 |  | Ensembl |
| 109 | L5 | NA | otherA |  | Arabian camel (*Camelus dromedarius*) L5 |  | Ensembl |
| 110 | L6 | NA | otherA |  | Asiatic black bear (*Ursus thibetanus thibetanus*) L6 |  | Ensembl |
| 111 | R1 | NA | otherA |  | Abingdon island giant tortoise (*Chelonoidis abingdonii*) R1 |  | Ensembl |
| 112 | R2 | NA | otherA |  | Agassiz's desert tortoise (*Gopherus agassizii*) R2 |  | Ensembl |
| 113 | R3 | NA | otherA |  | Argentine black and white tegu (*Salvator merianae*) R3 |  | Ensembl |
| 114 | R4 | NA | otherA |  | Australian saltwater crocodile (*Crocodylus porosus*) R4 |  | Ensembl |
| 115 | R5 | NA | otherA |  | Blue-ringed sea krait (*Laticauda laticaudata*) R5 |  | Ensembl |
| 116 | R6 | NA | otherA |  | Central bearded dragon (*Pogona vitticeps*) R6 |  | Ensembl |
| 117 | x1 | NA | otherA |  | Armadillo (*Dasypus novemcinctus*) x1 |  | Ensembl |
| 118 | x2 | NA | otherA |  | Sloth (*Choloepus hoffmanni*) x2 |  | Ensembl |
| 119 | x3 | NA | otherA |  | Elephant (*Loxodonta africana*) x3 |  | Ensembl |
| 120 | x4 | NA | otherA |  | Hyrax (*Procavia capensis*) x4 |  | Ensembl |
| 121 | x5 | NA | otherA |  | Lesser hedgehog tenrec (*Echinops telfairi*) x5 |  | Ensembl |
| 122 | x6 | NA | otherA |  | Leishan spiny toad (*Leptobranchium leishanense*) x6 |  | Ensembl |
| 123 | p1 | NA | primates |  | p1 *Aotus nancymae*.Anan_2.0.dna.toplevel |  | Ensembl |
| 124 | p10 | NA | primates |  | p10 *Papio anubis*.Panubis1.0.dna.toplevel |  | Ensembl |
| 125 | p11 | NA | primates |  | p11 *Ptilocolobus tephrosceles* ASM277652v2.dna.toplevel |  | Ensembl |
| 126 | p12 | NA | primates |  | p12 *Pongo abelii*.Susie_PABv2.dna.toplevel |  | Ensembl |
| 127 | p13 | NA | primates |  | p13 *Colobus angolensis palliatus*.Cang.pa_1.0.dna.toplevel |  | Ensembl |
| 128 | p14 | NA | primates |  | p14 *Rhinopithecus bieti*.ASM169854v1.dna.toplevel |  | Ensembl |
| 129 | p15 | NA | primates |  | p15 *Saimiri boliviensis boliviensis*.SaiBol1.0.dna.toplevel |  | Ensembl |
| 130 | p16 | NA | primates |  | p16 *Otolemur garnettii*.OtoGar3.dna.toplevel |  | Ensembl |
| 131 | p17 | NA | primates |  | p17 *Propithecus coquereli*.Pcoq_1.0.dna.toplevel |  | Ensembl |
| 132 | p18 | NA | primates |  | p18 *Macaca fascicularis*.Macaca_fascicularis_6.0.dna.toplevel |  | Ensembl |
| 133 | p19 | NA | primates |  | p19 *Mandrillus leucophaeus*.Mleu.le_1.0.dna.toplevel |  | Ensembl |
| 134 | p2 | NA | primates |  | p2 *Callithrix jacchus*.mCalJac1.pat.X.dna.toplevel |  | Ensembl |
| 135 | p20 | NA | primates |  | p20 *Theropithecus gelada*.Tgel_1.0.dna.toplevel |  | Ensembl |
| 136 | p21 | NA | primates |  | p21 *Nomascus leucogenys*.Nleu_3.0.dna.toplevel |  | Ensembl |
| 137 | p22 | NA | primates |  | p22 *Prolimur simus*.Prosim_1.0.dna.toplevel |  | Ensembl |
| 138 | p23 | NA | primates |  | p23 *Macaca mulatta*.Mmul_10.dna.toplevel |  | Ensembl |
| 139 | p24 | NA | primates |  | p24 *Aotus nancymae*.Anan_2.0.dna.toplevel |  | Ensembl |
| 140 | p25 | NA | primates |  | p25 *Microcebus murinus*.Mmur_3.0.dna.toplevel |  | Ensembl |
| 141 | p26 | NA | primates |  | p26 *Papio anubis*.Panubis1.0.dna.toplevel |  | Ensembl |
| 142 | p27 | NA | primates |  | p27 *Cebus imitator*.Cebus_imitator-1.0.dna.toplevel |  | Ensembl |
| 143 | p28 | NA | primates |  | p28 *Macaca nemestrina*.Mnem_1.0.dna.toplevel |  | Ensembl |
| 144 | p29 | NA | primates |  | p29 *Cercopithecus atys*.Caty_1.0.dna.toplevel |  | Ensembl |
| 145 | p3 | NA | primates |  | p3 *Cercopithecus atys*.Caty_1.0.dna.toplevel |  | Ensembl |
| 146 | p30 | NA | primates |  | p30 *Carlito syrichta*.Tarsyich-2.0.1.dna.toplevel |  | Ensembl |
| 147 | p31 | NA | primates |  | p31 *Ptilocolobus tephrosceles* ASM277652v2.dna.toplevel |  | Ensembl |
| 148 | p32 | NA | primates |  | p32 *Chlorocebus sabaeus*.ChlSab1.1.dna.toplevel |  | Ensembl |
| 149 | p33 | NA | primates |  | p33 *Callithrix jacchus*.mCalJac1.pat.X.dna.toplevel |  | Ensembl |
| 150 | p4 | NA | primates |  | p4 *Chlorocebus sabaeus*.ChlSab1.1.dna.toplevel |  | Ensembl |
| 151 | p5 | NA | primates |  | p5 *Gorilla gorilla*.gorGor4.dna.toplevel |  | Ensembl |
| 152 | p6 | NA | primates |  | p6 *Homo sapiens*.GRCh38.dna.toplevel |  | Ensembl |
| 153 | p7 | NA | primates |  | p7 *Macaca nemestrina*.Mnem_1.0.dna.toplevel |  | Ensembl |
| 154 | p8 | NA | primates |  | p8 *Pan paniscus*.panpan1.1.dna.toplevel |  | Ensembl |
| 155 | p9 | NA | primates |  | p9 *Pan troglodytes*.Pan_tro_3.0.dna.toplevel |  | Ensembl |
| 156 | bz1 | Brazil (a) | SouthAm | 8000 | Sumidouro Cave, Lagoa Santa Brazil | 41 | PRJEB29074 |
| 157 | ch1 | Chile(a) | SouthAm | 4700 | Ayayema | 35 | PRJEB29074 |
| 158 | me1 | Mexica | SouthAm |  | Mexican LosAngeL-1 | 103 | PRJEB31736 |
| 159 | ur1 | Uruguay(a) | SouthAm | 668 | Uruguay (CH13) | 19 | PRJEB48360 |
| 160 | ur2 | Uruguay(a) | SouthAm | 1400 | Uruguay (CH198) | 12 | PRJEB48360 |
| 161 | ap1 | China | otherB |  | SRR1552607 Alpaca genome sequencing 羊驼 | 45 | PRJNA233565 |
| 162 | bc1 | NA | otherB |  | Border Collie-1 | 63 | PRJEB4544 |
| 163 | bc2 | NA | otherB |  | Border Collie-2 | 97 | PRJEB36029 |
| 164 | cm1 | IRAN | otherB |  | CAMELcm1- IRAN_B_Run_189 | 95 | SRP107089 |
| 165 | cm2 | NA | otherB |  | Camel-2 | 80 | SRP107089 |
| 166 | dp1 | NA | otherB |  | Atlantic bottlenose dolphin (*Tursiops truncatus*) | 89 | PRJNA476133 |
| 167 | dp3 | NA | otherB |  | Atlantic bottlenose dolphin (*Tursiops truncatus*) | 104 | PRJNA20367 |
| 168 | ha1 | NA | otherB |  | USA Hawaiian SAMEA3302908 | 106 | PRJEB9586 |
| 169 | hu1 | NA | otherB |  | humpback whale | 9 | GCA_004329385 |
| 170 | pr1 | Australia | otherB |  | Parrot,male budgerigar (*Melopsittacus undulatus*) | 89 | PRJEB1588 |

Note: (a): ancient sample; SouthAm: South America; NorthAm: North America. References are ENA or SRA accession numbers plus Ensembl (fa or fna formats with the file size of around 200M~50G) . NA: not applicable. OtherA and Other B are two groups of samples mainly for animals.

### Language/Cognition genes and their SNPs

Language is an emergent complicated function of human being, though many other animals also have their own ‘languages’. If a gene mutation is statistically or experimentally associated with a certain language function loss, it would be called language gene. For both language gene and cognition gene, SNP sites in the dbSNP database were selected in a way that the each whole gene region was relatively equally spanned by the selected sites, plus those already with known clinical effects (seen in the Genecards database). Table 2 listed 36 language/cognition genes, and a total 239 SNPs from 18 language genes were selected for this study (Table 3), while 223 SNPs from 18 cognition genes were selected (Table 4).

**Table 2.** Selected language/Cognition genes in this study ^[9-15]^.

|  | Gene | Language gene | Cognition gene | Function or Compromised ability (example) when mutated |
| --- | --- | --- | --- | --- |
| 1 | FOXP1 | ✓ |  | Expressive language |
| 2 | FOXP2 | ✓ |  | Speech |
| 3 | CNTNAP2 | ✓ |  | Early language development |
| 4 | RBFOX2 | ✓ |  | Reading, language |
| 5 | TPK1 | ✓ |  | Syntactic and lexical ability |
| 6 | DCDC2 | ✓ |  | Reading, dyslexia |
| 7 | KIAA0319 | ✓ |  | Reading, dyslexia |
| 8 | TM4SF20 | ✓ |  | Language delay; communication disorder |
| 9 | FLNC | ✓ |  | Reading, language |
| 10 | ATP2C2 | ✓ |  | Memory |
| 11 | ROBO1 | ✓ |  | Phonological buffer |
| 12 | ROBO2 | ✓ |  | Expressive vocabulary |
| 13 | CMIP | ✓ |  | Reading, memory |
| 14 | DYX1C1 | ✓ |  | Reading, dyslexia |
| 15 | NFXL1 | ✓ |  | Speech |
| 16 | SRGAP2 | ✓ |  | vocal learning, vital for cortical neuron development |
| 17 | ARHGAP11B |  | ✓ | Hominin-specific protein that promotes development and evolutionary expansion of the brain neocortex |
| 18 | ASPM | ✓ | ✓ | ASPM (Assembly Factor For Spindle Microtubules) is a protein coding gene. Diseases associated with ASPM include Microcephaly, Primary, Autosomal Recessive and Primary Autosomal Recessive Microcephaly. |
| 19 | MCPH1 | ✓ | ✓ | A protein coding gene. Diseases associated with MCPH1 include Microcephaly, Primary, Autosomal Recessive and Lymphatic Malformation; |
| 20 | CHRM2 |  | ✓ | A nervous system gene associated with depression disorder |
| 21 | IGF2R |  | ✓ | Insulin-like growth factor gene associated with behavior/neurological phenotype |
| 22 | DTNBP1 |  | ✓ | A protein coding gene likely associated with schizophrenia |
| 23 | Snap25 |  | ✓ | A gene associated with neurotransmitter release |
| 24 | Fads2 |  | ✓ | A member of the fatty acid desaturase, associated with craniofacial abnormalities |
| 25 | Dab1 |  | ✓ | A gene linked with nervous system development |
| 26 | NBPF8 |  | ✓ | A gene associated with microcephaly, macrocephaly, autism, schizophrenia, cognitive disability |
| 27 | HAR1A |  | ✓ | A gene whose expression levels associated with memory and cognitive abilities |
| 28 | GNB5 |  | ✓ | G Protein Subunit Beta 5 associated with Intellectual Developmental Disorder, Language Delay And Attention Deficit-Hyperactivity Disorder/Cognitive Impairment |
| 29 | NRXN1 |  | ✓ | Neurexin 1 required for efficient neurotransmission and formation of synaptic contacts |
| 30 | DCC |  | ✓ | DCC Netrin 1 Receptor, a gene mediates axon guidance of neuronal growth cones and associated with Impaired Intellectual Development |
| 31 | GRID2 |  | ✓ | Glutamate Ionotropic Receptor Delta Type Subunit 2, predominant excitatory neurotransmitter receptors in the mammalian brain |
| 32 | EP300 |  | ✓ | E1A Binding Protein P300, associated with rare neurological diseases and impairment of intellectual development |
| 33 | KMT2D |  | ✓ | Lysine Methyltransferase 2D, associated with intellectual disability and eye diseases |
| 34 | NOTCH2NL |  | ✓ | A gene promotes neural progenitor proliferation and evolutionary expansion of the brain neocortex |
| 35 | THSD7B |  | ✓ | Associated with eye diseases/neuronal diseases |
| 36 | CASC5 |  | ✓ | Potentially associated with brain size of East Asian |

**Table 3.** Tested 239 SNPs of 18 language genes.

| snp code | snp info | snp code | snp info | snp code | snp info |
| --- | --- | --- | --- | --- | --- |
| ASPM10 | ASPM rs10754215 | FOXP1-2 | FOXP1 rs75214049 | ROBO1-4 | ROBO1 rs35456279 |
| ASPM-1 | ASPM rs12677 | FOXP1-1 | FOXP1 rs76145927 | ROBO1-3 | ROBO1 rs6795556 |
| ASPM-5 | ASPM rs1332663 | FXP1 | FOXP1 rs7638391 | ROBO1-2 | ROBO1 rs77350918 |
| ASPM-6 | ASPM rs1571964 | FOXP1-6 | FOXP1 rs7639736 | ROBO1-8 | ROBO1-rs1378638 |
| ASPM-7 | ASPM rs2878749 | FOXP1-16 | FOXP1-rs1288693 | ROBO1-6 | ROBO1-rs162423 |
| ASPM-8 | ASPM rs3737110 | FOXP1-18 | FOXP1-rs1463951 | ROBO1-7 | ROBO1-rs331168 |
| ASPM-9 | ASPM rs4915337 | FOXP1-17 | FOXP1-rs1733518 | ROBO1-13 | ROBO1-rs3923148 |
| ASPM-2 | ASPM rs877897 | FOXP1-13 | FOXP1-rs17803583 | ROBO1-11 | ROBO1-rs4130219 |
| ASPM-3 | ASPM rs955927 | FOXP1-12 | FOXP1-rs200643313 | ROBO1-12 | ROBO1-rs4130431 |
| ASPM-4 | ASPM rs964201 | FOXP1-14 | FOXP1-rs2044341412 | ROBO1-9 | ROBO1-rs716681 |
| ATP-8 | ATP2C2 rs13334642 | FOXP1-19 | FOXP1-rs2048059 | ROBO1-5 | ROBO1-rs80030397 |
| ATP-7 | ATP2C2 rs16973859 | FOXP1-15 | FOXP1-rs722261 | ROBO1-10 | ROBO1-rs991787 |
| ATP-11 | ATP2C2 rs2435172 | FOXP2-1 | FOXP2 rs10227893 | ROBO2-9 | ROBO2 rs1031377 |
| ATP-13 | ATP2C2 rs247818 | FOXP2-2 | FOXP2 rs10244649 | ROBO2-2 | ROBO2 rs10865561 |
| ATP-12 | ATP2C2 rs247885 | FOXP2-8 | FOXP2 rs1058335 | ROBO2-1 | ROBO2 rs11127602 |
| ATP-10 | ATP2C2 rs4782948 | FOXP2-22 | FOXP2 rs114972925 | ROBO2-8 | ROBO2 rs1163748 |
| ATP-9 | ATP2C2 rs4782970 | FOXP2-26 | FOXP2 rs115978361 | ROBO2-7 | ROBO2 rs1163749 |
| ATP-6 | ATP2C2 rs62050917 | FOXP2-27 | FOXP2 rs116557180 | ROBO2-6 | ROBO2 rs1163750 |
| ATP-5 | ATP2C2 rs62640931 | FOXP2-3 | FOXP2 rs12705977 | ROBO2-12 | ROBO2 rs144468527 |
| ATP-4 | ATP2C2 rs62640932 | FOXP2-10 | FOXP2 rs144807019 | ROBO2-13 | ROBO2 rs17525412 |
| ATP-3 | ATP2C2 rs62640935 | FOXP2-11 | FOXP2 rs182138317 | ROBO2-5 | ROBO2 rs3923744 |
| ATP-2 | ATP2C2 rs74038217 | FOXP2-23 | FOXP2 rs191654848 | ROBO2-4 | ROBO2 rs3923745 |
| ATP-1 | ATP2C2 rs78371901 | FOXP2-24 | FOXP2 rs560859215 | ROBO2-3 | ROBO2 rs5788280 |
| CMI-4 | CMIP rs114894868 | FOXP2-30 | FOXP2 rs563023653 | ROBO2-11 | ROBO2 rs78817248 |
| CMI-13 | CMIP rs1187121850 | FOXP2-25 | FOXP2 rs577428580 | ROBO2-14 | ROBO2-rs12171318 |
| CMI-11 | CMIP rs16955675 | FOXP2-4 | FOXP2 rs61732741 | ROBO2-19 | ROBO2-rs1372422 |
| CMI-3 | CMIP rs183075361 | FOXP2-9 | FOXP2 rs61753357 | ROBO2-20 | ROBO2-rs1372427 |
| CMI-2 | CMIP rs183876152 | FOXP2-5 | FOXP2 rs61758964 | ROBO2-21 | ROBO2-rs1503125 |
| CMI-1 | CMIP rs201316817 | FOXP2-6 | FOXP2 rs62640396 | ROBO2-16 | ROBO2-rs17203 |
| CMI-12 | CMIP rs2288011 | FOXP2-7 | FOXP2 rs73210755 | ROBO2-15 | ROBO2-rs264546 |
| CMI-10 | CMIP rs34119643 | FOXP2-29 | FOXP2 rs7782412 | ROBO2-17 | ROBO2-rs699456 |
| CMI-9 | CMIP rs35429777 | FOXP2-28 | FOXP2 rs7795372 | ROBO2-18 | ROBO2-rs873596 |
| CMI-8 | CMIP rs57603843 | FOXP2-31 | FOXP2 rs7799652 | SRGAP2 | SRGAP2 rs1350526469 |
| CMI-7 | CMIP rs60152409 | FOXP2-14 | FOXP2-rs531957198 | SRGAP2 | SRGAP2 rs1361269 |
| CMI-6 | CMIP rs74031247 | FOXP2-16 | FOXP2-rs718378 | SRGAP2 | SRGAP2 rs1476372 |
| CMI-5 | CMIP rs79979027 | FOXP2-17 | FOXP2-rs724419 | SRGAP2 | SRGAP2 rs17018890 |
| CNTN-6 | CNTNAP2 rs1062071 | FOXP2-12 | FOXP2-rs747126499 | SRGAP2 | SRGAP2 rs1754475 |
| CNTN-5 | CNTNAP2 rs1062072 | FOXP2-15 | FOXP2-rs773664240 | SRGAP2 | SRGAP2 rs2244510 |
| CNTN-4 | CNTNAP2 rs1468370 | FOXP2-18 | FOXP2-rs776920 | SRGAP2 | SRGAP2 rs2987927 |
| CNTN-3 | CNTNAP2 rs1479837 | FOXP2-19 | FOXP2-rs814066 | SRGAP2 | SRGAP2 rs502336 |
| CNTN-2 | CNTNAP2 rs1637841 | FOXP2-21 | FOXP2-rs940468 | SRGAP2 | SRGAP2 rs508058 |
| CNTN-1 | CNTNAP2 rs1637842 | FOXP2-20 | FOXP2-rs956016 | SRGAP2 | SRGAP2 rs523647 |
| CNTN-12 | CNTNAP2 rs2373284 | KIA-8 | KIAA0319 rs10946705 | SRGAP2C | SRGAP2C-rs1546945 |
| CNTN-10 | CNTNAP2 rs3194 | KIA-3 | KIAA0319 rs114195393 | SRGAP2C | SRGAP2C-rs1769152 |
| CNTN-11 | CNTNAP2 rs535454043 | KIA-12 | KIAA0319 rs115399701 | SRGAP2C | SRGAP2C-rs2993869 |
| CNTN-13 | CNTNAP2 rs61732853 | KIA-2 | KIAA0319 rs117692893 | SRGAP2C | SRGAP2C-rs458475 |
| CNTN-9 | CNTNAP2 rs700308 | KIA-1 | KIAA0319 rs138160539 | SRGAP2C | SRGAP2C-rs493257 |
| CNTN-8 | CNTNAP2 rs700309 | KIA-11 | KIAA0319 rs150584710 | SRGAP2C | SRGAP2C-rs499258 |
| CNTN-7 | CNTNAP2 rs987456 | KIA-4 | KIAA0319 rs699461 | SRGAP2C | SRGAP2C-rs519348 |
| D CD-12 | D CDC2 rs190254728 | KIA-5 | KIAA0319 rs699462 | SRGAP2C | SRGAP2C-rs545109 |
| D CD-2 | D CDC2 rs2274305 | KIA-6 | KIAA0319 rs699463 | SRGAP2C | SRGAP2C-rs561290 |
| D CD-5 | D CDC2 rs33914824 | KIA-7 | KIAA0319 rs730860 | SRGAP2C | SRGAP2C-rs71251644 |
| D CD-11 | D CDC2 rs33943110 | KIA-9 | KIAA0319 rs75674723 | TM8 | TM4SF20 rs13415654 |
| D CD-3 | D CDC2 rs34584835 | KIA-10 | KIAA0319 rs75720688 | TM10 | TM4SF20 rs137891000 |
| D CD-1 | D CDC2 rs35029429 | KIA-13 | KIAA0319 rs7770041 | TM7 | TM4SF20 rs4408717 |
| D CD-9 | D CDC2 rs3789219 | MCPH1-3 | MCPH1 rs1057091 | TM6 | TM4SF20 rs4428010 |
| D CD-8 | D CDC2 rs3846827 | MCPH1-8 | MCPH1 rs115556798 | TM5 | TM4SF20 rs4438464 |
| D CD-7 | D CDC2 rs9460973 | MCPH1-4 | MCPH1 rs1550689 | TM2 | TM4SF20 rs44675173 |
| D CD-6 | D CDC2 rs9467075 | MCPH1-5 | MCPH1 rs1550691 | TM4 | TM4SF20 rs4673192 |
| FLN-11 | FLNC rs117864464 | MCPH1-6 | MCPH1 rs1550696 | TM3 | TM4SF20 rs4675172 |
| FLN-10 | FLNC rs2249128 | MCPH1-7 | MCPH1 rs1961222 | TM1 | TM4SF20 rs6724955 |
| FLN-9 | FLNC rs2291558 | MCPH1-1 | MCPH1 rs2583 | TM9 | TM4SF20 rs80305648 |
| FLN-8 | FLNC rs2291560 | MCPH1 -9 | MCPH1 rs7814961 | TM4SF20-14 | TM4SF20-rs10168278 |
| FLN-7 | FLNC rs2291561 | MCPH1 -2 | MCPH1 rs890223 | TM4SF20-15 | TM4SF20-rs4675173 |
| FLN-6 | FLNC rs2291562 | MCPH1-10 | MCPH1 rs895973 | TM4SF20-11 | TM4SF20-rs7568026 |
| FLN-5 | FLNC rs2291563 | NFX-8 | NFXL1 rs1036681 | TM4SF20-12 | TM4SF20-rs7574414 |
| FLN-4 | FLNC rs2291565 | NFXL1-14 | NFXL1 rs12651301 | TM4SF20-13 | TM4SF20-rs9678000 |
| FLN-3 | FLNC rs2291566 | NFX-12 | NFXL1 rs13152765 | TPK-1 | TPK1 rs113536847 |
| FLN-2 | FLNC rs2291568 | NFX-7 | NFXL1 rs1371730 | TPK-6 | TPK1 rs12333969 |
| FLN-1 | FLNC rs2291569 | NFX-6 | NFXL1 rs1440228 | TPK-5 | TPK1 rs17170295 |
| FLN-12 | FLNC rs35281128 | NFX-11 | NFXL1 rs147017712 | TPK-4 | TPK1 rs28380423 |
| FLN-13 | FLNC rs371111092 | NFX-5 | NFXL1 rs1545200 | TPK-9 | TPK1 rs67644764 |
| FOXP1-8 | FOXP1 rs1053797 | NFX-4 | NFXL1 rs1812964 | TPK-7 | TPK1 rs6953807 |
| FOXP1-5 | FOXP1 rs11914627 | NFX-3 | NFXL1 rs1822029 | TPK-3 | TPK1 rs77358162 |
| FOXP1-9 | FOXP1 rs144080925 | NFX-2 | NFXL1 rs1822030 | TPK-2 | TPK1 rs79464600 |
| FOXP1-11 | FOXP1 rs147756430 | NFX-1 | NFXL1 rs1964425 | TPK1-13 | TPK1-rs228582 |
| FOXP1-7 | FOXP1 rs1499893 | NFX-13 | NFXL1 rs34323060 | TPK1-11 | TPK1-rs38045 |
| FOXP1-4 | FOXP1 rs17008063 | NFX-10 | NFXL1 rs920462 | TPK1-12 | TPK1-rs38046 |
| FOXP1-10 | FOXP1 rs17008224 | NFX-9 | NFXL1 rs978094 | TPK1-10 | TPK1-rs41239 |
| FOXP1-3 | FOXP1 rs17008544 | ROBO1-1 | ROBO1 rs34841026 |  |  |

**Table 4.** Tested 223 SNPs of 18 cognition genes.

| snp code | snp info | snp code | snp info | snp code | snp info |
| --- | --- | --- | --- | --- | --- |
| NRXN1-1 | NRXN1 rs201544418 | DCC-5 | DCC rs141716650 | KMT2D-13 | KMT2D rs1943280408 |
| NRXN1-2 | NRXN1 rs886056179 | DCC-6 | DCC rs1057518248 | KMT2D-14 | KMT2D rs371223664 |
| NRXN1-3 | NRXN1 rs201539806 | DCC-7 | DCC rs141813053 | KMT2D-15 | KMT2D rs112170602 |
| NRXN1-4 | NRXN1 rs772333323 | DCC-8 | DCC rs1085307773 | KMT2D-16 | KMT2D rs1592118953 |
| NRXN1-5 | NRXN1 rs2303298 | DCC-9 | DCC rs775565634 | KMT2D-17 | KMT2D rs201119371 |
| NRXN1-6 | NRXN1 rs886056171 | DCC-10 | DCC rs387906555 | KMT2D-18 | KMT2D rs1555187117 |
| NRXN1-7 | NRXN1 rs750156118 | FADS2-1 | FADS2 rs1364380970 | KMT2D-19 | KMT2D rs1942683224 |
| NRXN1-8 | NRXN1 rs2091595186 | FADS2-2 | FADS2 rs2066989156 | KMT2D-20 | KMT2D rs1555184538 |
| NRXN1-9 | NRXN1 rs562219421 | FADS2-3 | FADS2 rs118041921 | MCPH1-01 | MCPH1 rs2920676 |
| NRXN1-10 | NRXN1 rs761776814 | FADS2-4 | FADS2 rs149777687 | MCPH1-02 | MCPH1 rs2305023 |
| NRXN1-11 | NRXN1 rs760815320 | FADS2-5 | FADS2 rs1453175607 | MCPH1-03 | MCPH1 rs1550697 |
| NRXN1-12 | NRXN1 rs541005670 | FADS2-6 | FADS2 rs1591161560 | MCPH1-04 | MCPH1 rs587783733 |
| THSD7B-1 | THSD7B rs91435 | FADS2-7 | FADS2 rs1447645723 | MCPH1-05 | MCPH1 rs775942126 |
| THSD7B-2 | THSD7B rs35967139 | FADS2-8 | FADS2 rs2135964926 | MCPH1-06 | MCPH1 rs201721894 |
| THSD7B-3 | THSD7B rs114612136 | FADS2-9 | FADS2 rs2067344862 | MCPH1-07 | MCPH1 rs2053618 |
| THSD7B-4 | THSD7B rs149172693 | FADS2-10 | FADS2 rs972367375 | MCPH1-08 | MCPH1 rs145820898 |
| THSD7B-5 | THSD7B rs189224302 | FADS2-11 | FADS2 rs1168297160 | MCPH1-09 | MCPH1 rs548329168 |
| THSD7B-6 | THSD7B rs373333594 | GNB5-1 | GNB5 rs190432484 | MCPH1-010 | MCPH1 rs115033462 |
| THSD7B-7 | THSD7B rs534053420 | GNB5-2 | GNB5 rs6493537 | MCPH1-011 | MCPH1 rs370275760 |
| THSD7B-8 | THSD7B rs962699493 | GNB5-3 | GNB5 rs113335851 | MCPH1-012 | MCPH1 rs2936531 |
| THSD7B-9 | THSD7B rs1162258857 | GNB5-4 | GNB5 rs766151886 | MCPH1-013 | MCPH1 rs2011423 |
| THSD7B-10 | THSD7B rs1280704421 | GNB5-5 | GNB5 rs770868918 | NBPF8-1 | NBPF8 rs320820 |
| THSD7B-11 | THSD7B rs1398759851 | GNB5-6 | GNB5 rs756678877 | NBPF8-2 | NBPF8 rs1210092201 |
| THSD7B-12 | THSD7B rs1573714708 | GNB5-7 | GNB5 rs1330914161 | NBPF8-3 | NBPF8 rs1293062874 |
| ARHGAP11B-1 | ARHGAP11B rs1342963824 | GNB5-8 | GNB5 rs147993382 | NBPF8-4 | NBPF8 rs1378831317 |
| ARHGAP11B-2 | ARHGAP11B rs2060186953 | GNB5-9 | GNB5 rs1253307264 | NBPF8-5 | NBPF8 rs1469790408 |
| ARHGAP11B-3 | ARHGAP11B rs1473937662 | GNB5-10 | GNB5 rs1452240112 | NBPF8-6 | NBPF8 rs1660411467 |
| ARHGAP11B-4 | ARHGAP11B rs921027344 | GNB5-11 | GNB5 rs2033629123 | NBPF8-7 | NBPF8 rs1660852853 |
| ARHGAP11B-5 | ARHGAP11B rs2140884449 | GNB5-12 | GNB5 rs372011977 | NBPF8-8 | NBPF8 rs1661374355 |
| ARHGAP11B-6 | ARHGAP11B rs1161395884 | GNB5-13 | GNB5 rs761399728 | NBPF8-9 | NBPF8 rs1661790471 |
| ARHGAP11B-7 | ARHGAP11B rs2060245516 | GRID2-1 | GRID2 rs80091080 | NBPF8-10 | NBPF8 rs2101536757 |
| ARHGAP11B-8 | ARHGAP11B rs374363097 | GRID2-2 | GRID2 rs115664626 | NBPF8-11 | NBPF8 rs518881 |
| ARHGAP11B-9 | ARHGAP11B rs942105401 | GRID2-3 | GRID2 rs78407646 | NBPF8-12 | NBPF8 rs1638305116 |
| ARHGAP11B-10 | ARHGAP11B rs1021221929 | GRID2-4 | GRID2 rs181918786 | NOTCH2NLA-1 | NOTCH2NLA rs8002 |
| ARHGAP11B-11 | ARHGAP11B rs2060352325 | GRID2-5 | GRID2 rs75225211 | NOTCH2NLA-2 | NOTCH2NLA rs868975060 |
| ARHGAP11B-12 | ARHGAP11B rs2060373053 | GRID2-6 | GRID2 rs142012040 | NOTCH2NLA-3 | NOTCH2NLA rs1299634089 |
| ASPM-01 | ASPM rs759485449 | GRID2-7 | GRID2 rs182933054 | NOTCH2NLA-4 | NOTCH2NLA rs1450495295 |
| ASPM-02 | ASPM rs1451306414 | GRID2-8 | GRID2 rs759075553 | NOTCH2NLA-5 | NOTCH2NLA rs1553806412 |
| ASPM-03 | ASPM rs774143329 | GRID2-9 | GRID2 rs150053332 | NOTCH2NLA-6 | NOTCH2NLA rs1553814256 |
| ASPM-04 | ASPM rs77424753 | GRID2-10 | GRID2 rs1728640535 | NOTCH2NLA-7 | NOTCH2NLA rs1571267596 |
| ASPM-05 | ASPM rs587783211 | GRID2-11 | GRID2 rs750331613 | NOTCH2NLA-8 | NOTCH2NLA rs1661787802 |
| ASPM-06 | ASPM rs587783215 | HAR1A-1 | HAR1A rs956279328 | NOTCH2NLA-9 | NOTCH2NLA rs1663242974 |
| ASPM-07 | ASPM rs41265225 | HAR1A-2 | HAR1A rs2066138086 | NOTCH2NLA-10 | NOTCH2NLA rs2102289207 |
| ASPM-08 | ASPM rs199422171 | HAR1A-3 | HAR1A rs1601177451 | NOTCH2NLA-11 | NOTCH2NLA rs1553818207 |
| ASPM-09 | ASPM rs199422181 | HAR1A-4 | HAR1A rs2122871899 | SNAP25-1 | SNAP25 rs363050 |
| ASPM-010 | ASPM rs199422200 | HAR1A-5 | HAR1A rs1029593231 | SNAP25-2 | SNAP25 rs769950821 |
| CHRM2-1 | CHRM2 rs566459725 | HAR1A-6 | HAR1A rs2066137438 | SNAP25-3 | SNAP25 rs201770060 |
| CHRM2-2 | CHRM2 rs139124053 | HAR1A-7 | HAR1A rs574068890 | SNAP25-4 | SNAP25 rs371883444 |
| CHRM2-3 | CHRM2 rs79607027 | HAR1A-8 | HAR1A rs2066137022 | SNAP25-5 | SNAP25 rs763997141 |
| CHRM2-4 | CHRM2 rs1057524597 | HAR1A-9 | HAR1A rs762381080 | SNAP25-6 | SNAP25 rs797044873 |
| CHRM2-5 | CHRM2 rs324651 | HAR1A-10 | HAR1A rs933579923 | SNAP25-7 | SNAP25 rs1568623929 |
| CHRM2-6 | CHRM2 rs1805069679 | HAR1A-11 | HAR1A rs2122871565 | SNAP25-8 | SNAP25 rs1555794286 |
| CHRM2-7 | CHRM2 rs1805074915 | IGF2R-1 | IGF2R rs76130099 | SNAP25-9 | SNAP25 rs362998 |
| CHRM2-8 | CHRM2 rs1440850408 | IGF2R-2 | IGF2R rs76235629 | SNAP25-10 | SNAP25 rs533404025 |
| CHRM2-9 | CHRM2 rs141951417 | IGF2R-3 | IGF2R rs8191881 | SNAP25-11 | SNAP25 rs79020892 |
| CHRM2-10 | CHRM2 rs774760812 | IGF2R-4 | IGF2R rs8191808 | SNAP25-12 | SNAP25 rs200030321 |
| CHRM2-11 | CHRM2 rs76394680 | IGF2R-5 | IGF2R rs55987511 | SNAP25-13 | SNAP25 rs1871070779 |
| DAB1-1 | DAB1 rs114941053 | IGF2R-6 | IGF2R rs150809922 | EP300-1 | EP300 rs1601642812 |
| DAB1-2 | DAB1 rs183171115 | IGF2R-7 | IGF2R rs199651009 | EP300-2 | EP300 rs2059220495 |
| DAB1-3 | DAB1 rs34466938 | IGF2R-8 | IGF2R rs193920850 | EP300-3 | EP300 rs142673005 |
| DAB1-4 | DAB1 rs17117702 | IGF2R-9 | IGF2R rs756631085 | EP300-4 | EP300 rs144626200 |
| DAB1-5 | DAB1 rs75129043 | IGF2R-10 | IGF2R rs121434588 | EP300-5 | EP300 rs1057518002 |
| DAB1-6 | DAB1 rs145999889 | IGF2R-11 | IGF2R rs121434587 | EP300-6 | EP300 rs1555912107 |
| DAB1-7 | DAB1 rs199645763 | KMT2D-1 | KMT2D rs1342235871 | EP300-7 | EP300 rs755816596 |
| DAB1-8 | DAB1 rs1855377 | KMT2D-2 | KMT2D rs111266743 | EP300-8 | EP300 rs1601636833 |
| DAB1-9 | DAB1 rs769093224 | KMT2D-3 | KMT2D rs1592145879 | EP300-9 | EP300 rs6002271 |
| DAB1-10 | DAB1 rs1570632701 | KMT2D-4 | KMT2D rs1057520167 | EP300-10 | EP300 rs1464734494 |
| DAB1-11 | DAB1 rs532864586 | KMT2D-5 | KMT2D rs1555198921 | EP300-11 | EP300 rs1114167305 |
| DAB1-12 | DAB1 rs12404008 | KMT2D-6 | KMT2D rs1243381790 | EP300-12 | EP300 rs199773872 |
| DAB1-13 | DAB1 rs183171115 | KMT2D-7 | KMT2D rs398123747 | EP300-13 | EP300 rs2059047548 |
| DAB1-14 | DAB1 rs141647150 | KMT2D-8 | KMT2D rs141231056 | EP300-14 | EP300 rs774840930 |
| DCC-1 | DCC rs1057519054 | KMT2D-9 | KMT2D rs746084513 | EP300-15 | EP300 rs750740148 |
| DCC-2 | DCC rs797044551 | KMT2D-10 | KMT2D rs794727420 | EP300-16 | EP300 rs886057556 |
| DCC-3 | DCC rs116498325 | KMT2D-11 | KMT2D rs75783546 |  |  |
| DCC-4 | DCC rs200099519 | KMT2D-12 | KMT2D rs398123755 |  |  |

### Genome sequence analysis software development

The SNP (Single Nucleotide Polymorphism) loci finding software based on hash tables primarily processes biological whole-genome files and rapidly identifies SNP loci within the genome by utilizing a hash table-based search algorithm, obtaining the specific values of the mutated bases at these loci (A/C/G/T). The software is written in Python. Initially, it processes three different formats of whole-genome files—fastq, fna, and fa—based on their unique characteristics, extracting gene sequences and generating standard format files that include all lines containing only ATCGN bases. Subsequently, it reads the SNP file and stores the information for each locus into a hash table. The target sequence is converted into binary representation according to the corresponding rules of A-00, T-01, C-10, G-11, and then into decimal form to serve as keys in the hash table. For the standard files, the software reads sequences in groups based on the matching length and converts them into the corresponding decimal representation for comparison with the keys in the hash table. If a match is successful and the number of matched bases does not exceed the limit, the matched bases and location information are added to the corresponding value. This process is repeated until the end of the file. Upon completion of each file match, the results are tallied to determine the matched bases and their respective quantities for each SNP locus, and the findings are outputted. During use, the software can process multiple genome files in batches and impose restrictions on the matching length and the number of matches for a single SNP. After extensive validation, this software has shown a significant improvement in speed compared to conventional matching algorithms and other software based on KMP (Knuth–Morris–Pratt) improved algorithms.

### Sample SNP information abstraction and PCA analysis

The authors used 010Edit software to extract SNP information from genome files, but most SNP information were abstracted with hash07plus03 software. In all 170 genomes, the sizes mainly range from 200M to 120G. Genomes with fastq format but less than 10G were generally neglected or only used as a reference. Principal Component Analysis(PCA)was performed using R packages FactoMineR, factoextra, ggrepel and ggplot2. R codes for PCA can be seen in reference [16].

### SNP profile similarity measurement among samples

In order to compare the similarities in SNP profiles between the samples of each genome file, we used three suitable similarity calculation methods to conduct similarity analysis on the SNP site bases matched by all samples, and obtained the following results. Method 1: Levenshtein Distance algorithm [17]. This method measures the difference between bases at two sites by calculating the edit distance, that is, the minimum number of operations (insertions, deletions, or substitutions) required to convert bases at one site into bases at another site. We combine all The editing distance of the bases at the site is summed and divided by the maximum number of bases in the calculated sample to obtain the difference rate, which is converted into similarity. Method 2: Smith-Waterman algorithm [18]. This method uses dynamic programming to find the local optimal alignment between two sequences, that is, the subsequence with the highest similarity between the two sequences. We spliced the bases of all sites into a string, then used this algorithm to calculate the similarity score between the two strings, and then normalized the scores to obtain the similarity rate. Method 3: Needleman-Wunsch algorithm [19]. This method is similar to method 2, but this method adopts a global optimal alignment scheme, that is, considering the overall similarity of the two sequences instead of the local similarity. We also splice the bases of all sites into a string, and then use this algorithm to construct a score matrix to calculate the similarity score between the two strings, and then obtain the similarity rate. We applied these three methods to the calculation of similarity, and obtained the similarity between any two samples, as well as the similarity of all samples relative to the reference sample p6, and also drew a similarity change curve to intuitively demonstrate changing trends.

## Results and Discussion

The PCA analysis results (Figures 1-3) show that among all samples, various lower animals are situated at the farthest left position, followed closely by a batch of ancient human samples. The genomic sequences of all animals are whole-genome sequences, whereas many ancient human genome sequences are incomplete, such as c20 and nd10, which only have 1.7G and 200M respectively. Therefore, the clustering of these ancient human samples with lower animals can only indicate that certain SNP patterns in these ancient human samples belong to a more primitive state, corresponding to the initial evolutionary stage of ancient humans. Modern humans are mostly located at the farthest right position in the figures, so from left to right, it essentially reflects the evolutionary stages from low to high. Interestingly, samples corresponding to each evolutionary stage seem to simultaneously include origins from Asia, Africa, Europe, and the Americas. This suggests that new evolutionary populations generally had enough time to spread across nearly all continents; with the current sample size, it appears that European samples, especially Neanderthal samples, have the highest occurrence rate in the earliest evolutionary stages of language genes and cognitive genes.

**Fig1.**
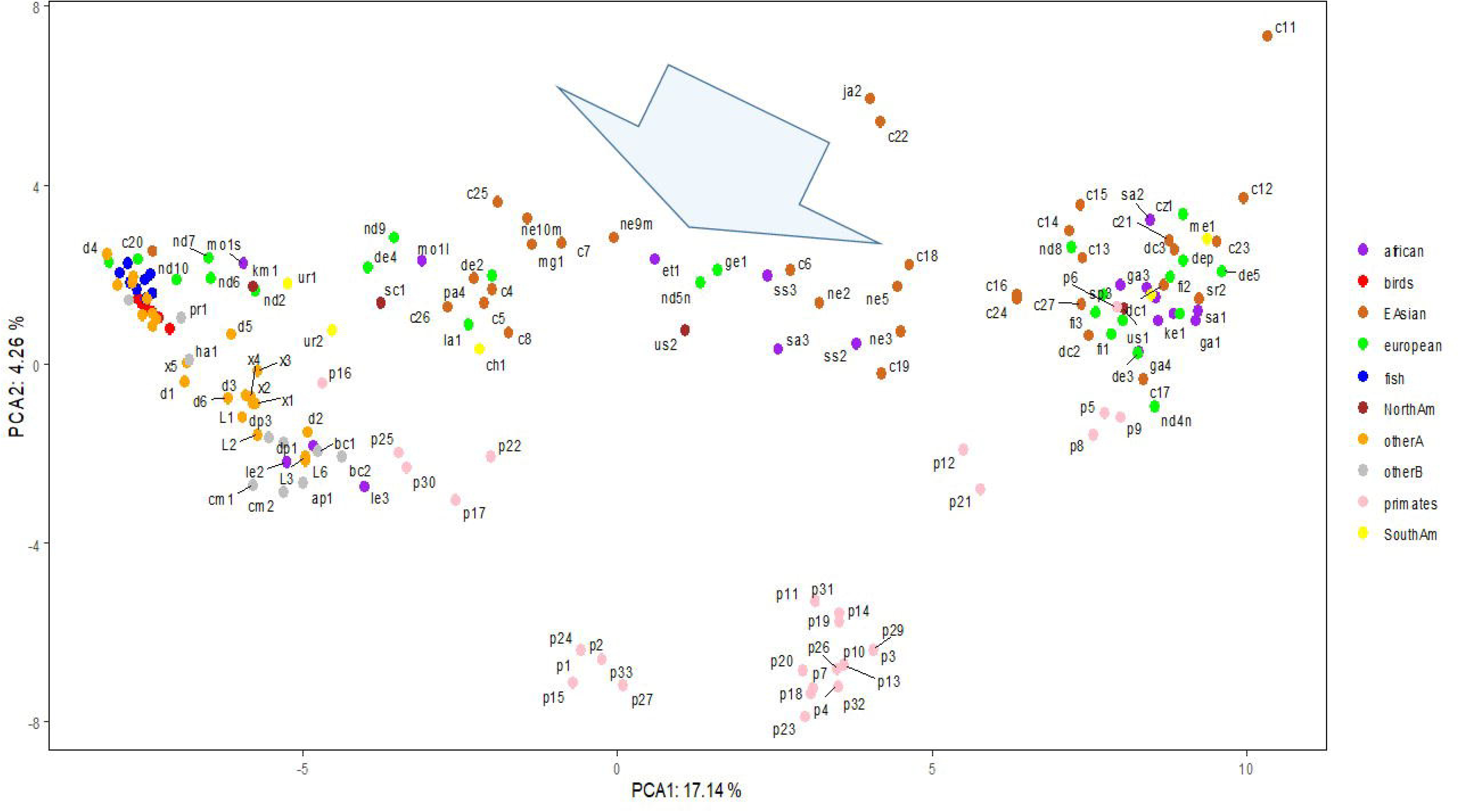
CG PCA result for SNPs in cognition genes.

The evolutionary process of language genes (Figure 2) differs significantly from that of cognitive genes (Figure 1). The early patterns of SNP diversity in language genes are very similar across all animal samples, whereas cognitive genes exhibit significant variation right from the start among the selected animal samples. There are two polymorphic states of cognitive genes at the beginning of evolution, one state is reflected in a group of intelligent animals (such as border collies, camels, alpacas, cetaceans, cheetahs, etc.) and primates, while the other state is directly reflected in certain animals and early human populations, and these two states gradually diverge and then come close again, but they never overlap; the closest distance between these two states is reflected in the orangutan species, because the polymorphic patterns of cognitive and language genes in chimpanzees and bonobos are closest to those of modern humans.

**Fig2.**
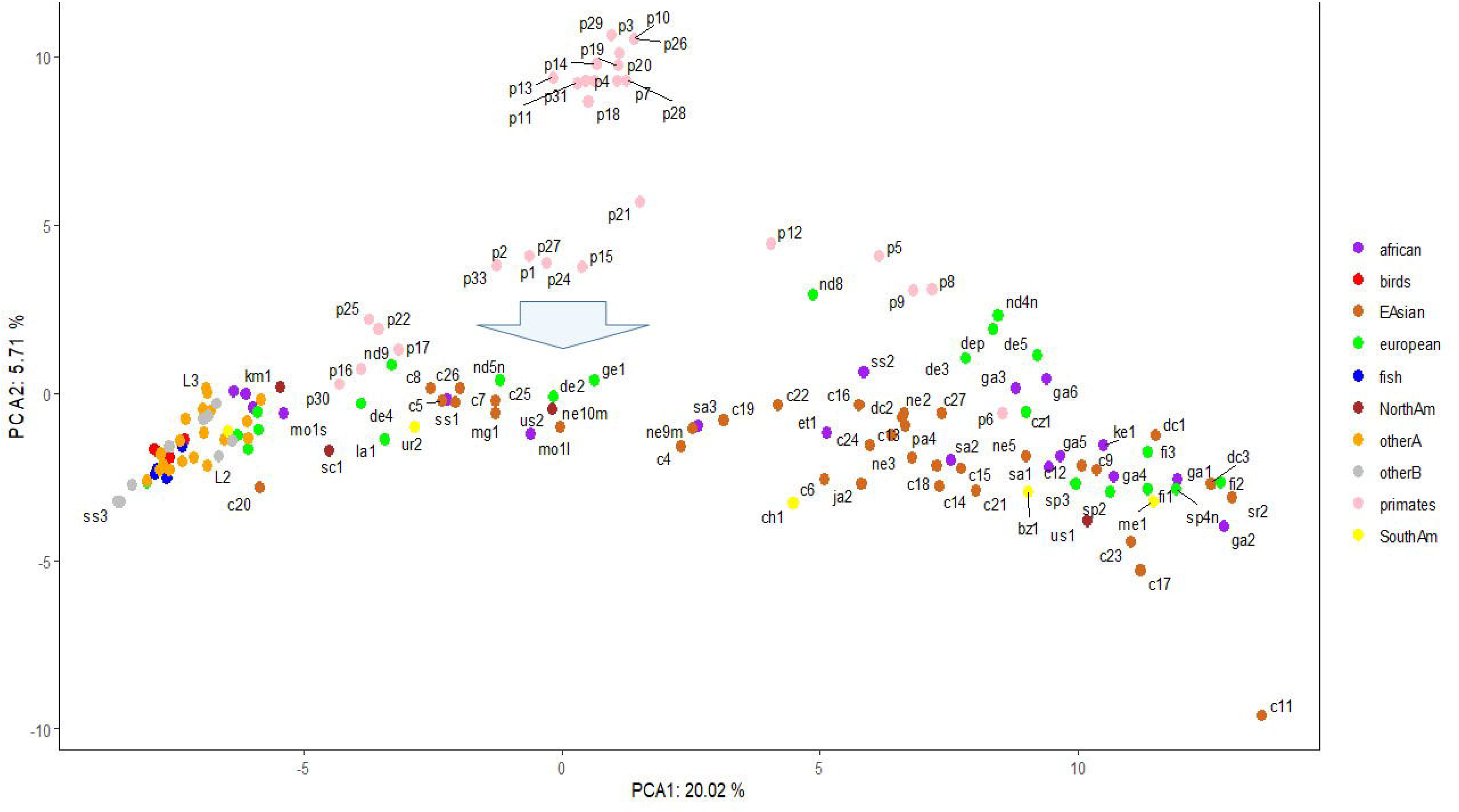
LG PCA result for SNPs in language genes.

**Fig3.**
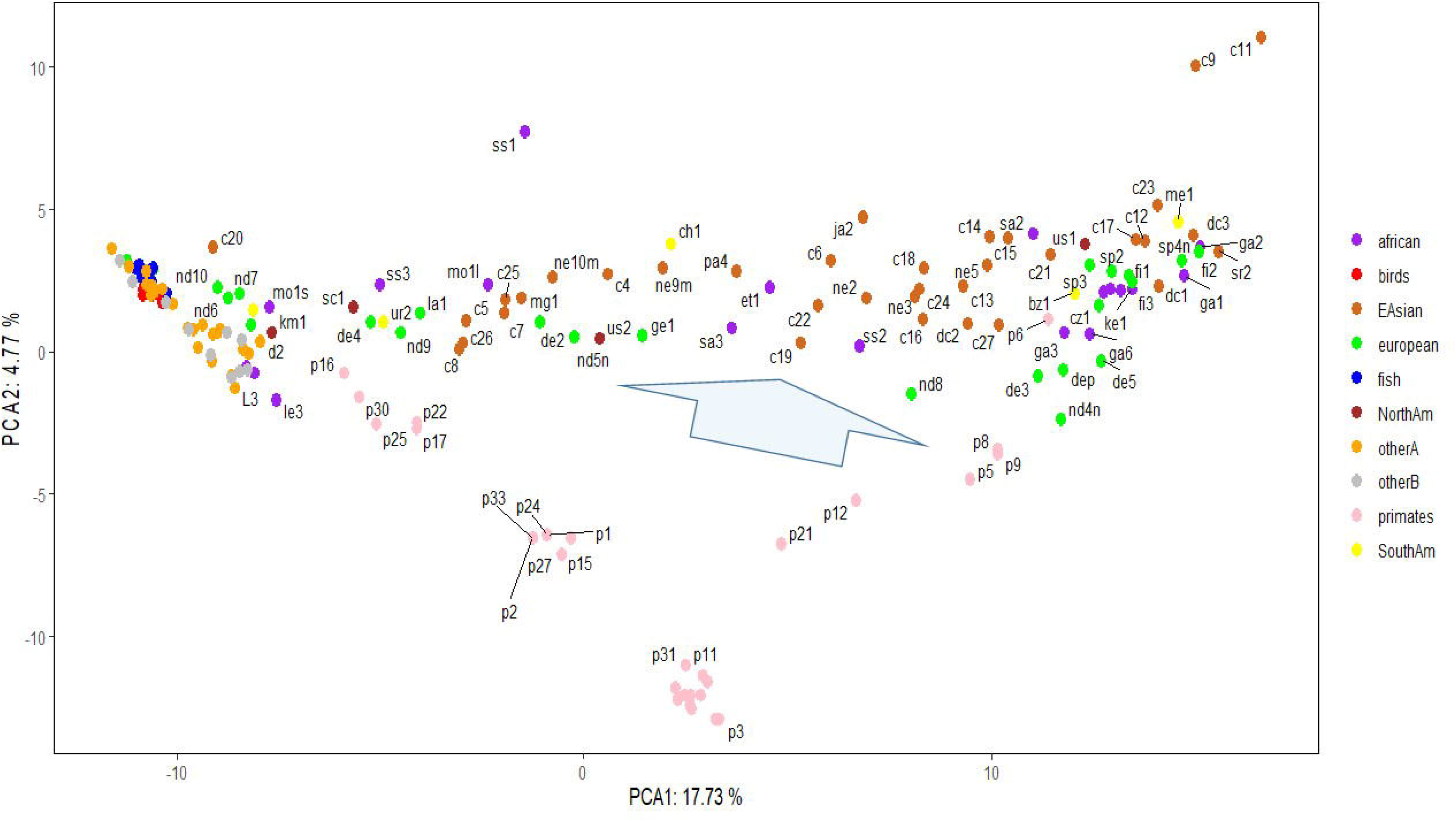
CGLG PCA result for SNPs in language and cognition genes.

The Asian sample c20 (Tianyuan) consistently appears at the earliest stage of evolution. Since the whole-genome sequence of this sample is not complete, the conclusions drawn from it are currently insufficient, but it generally tends to appear with European Neanderthals at the earliest stages of evolution, which at least suggests that some SNP loci of this sample may belong to the most ancient combination patterns. The Tianyuan genome possesses genomic features close to modern Asians, who carry approximately 4%–5% Neanderthal DNA shared by Upper Paleolithic Eurasians. The Tianyuan genome has a relatively closer relationship with present-day and ancient Asians than with Europeans [20]. European sample nd10 shares a similar situation with the Asian sample c20, that is, due to the extremely incomplete genome sequence, the results obtained lack sufficient credibility. The only reference worth mentioning is that some of its SNP patterns are in the oldest state of evolution. Clearly, from Fig 1-3, it is not feassible to discern whether the oldest SNP patterns originated from Asia, Africa, or Europe.

Comparing the evolutionary process of language genes and cognitive genes in European samples, the basic conclusion is that Neanderthals have consistently been at the early stages of evolution. Among all African samples, the Moroccan sample seems to be at the earliest stage of evolution, and indeed there is literature supporting that the earliest Homo sapiens from Africa originated from the Moroccan region [21-22];

Quasi-quantitative measurement among CG/LG/CGLG SNP profiles for 170 samples is essential to observe the potential points at which big changes happen. SNP profile similarity measurement results (Fig4) may tell something. In Fig4A, there are roughly two points (see two arrows) showing significant value changes for three different measurements (Levenshtein Distance, Smith-Waterman similarity and Needleman-Wunsch similarity; those with three measurements significantly changed at the same time can be regarded as turning points), and around the two points, there are total about 14 samples (ne9m, et1, nd5n, sa3, us2, ge1, ss3, ss2,c19, ne3,c16, ne2, c24,c15). All these 14 samples are located at the big arrow position in Fig1. In Fig4B, there are also roughly two turning points (see two arrows), around which there are basically 9 samples (c5, c26, c7, c8, mg1, nd5n, us2, ge1, et1). In Fig4C, there are another two turning points that harbor 5 samples (us2, ge1, ja1, c19 and nd8). The Fig4 demonstrated that CG and LG plus CGLG profiles have 2 common samples (ge1and us2) at their turning points, suggesting that Europe or North America might be also key sites for human evolution, at which sites some critical changes in both language development and cognition function especially simbolic thinking may experience an accumulative jump.

Interestingly, primate samples can be observed at the aforementioned turning points, and the extent of their presence varies across the turning points in Figs. 4A, 4B, and 4C. This also results in a noticeable difference in the number of human samples at these turning points, which are 14, 9, and 5 respectively. Detailed information can be found in Supplementary files 1 to 3. The primate samples that appear at the turning point in Fig. 4A are p33, p24, p20, p27, and p15; those at the turning point in Fig. 4B are p2, p33, and p22; and those at the turning point in Fig. 4C are p1, p26, p24, and p13.

**Fig4.**
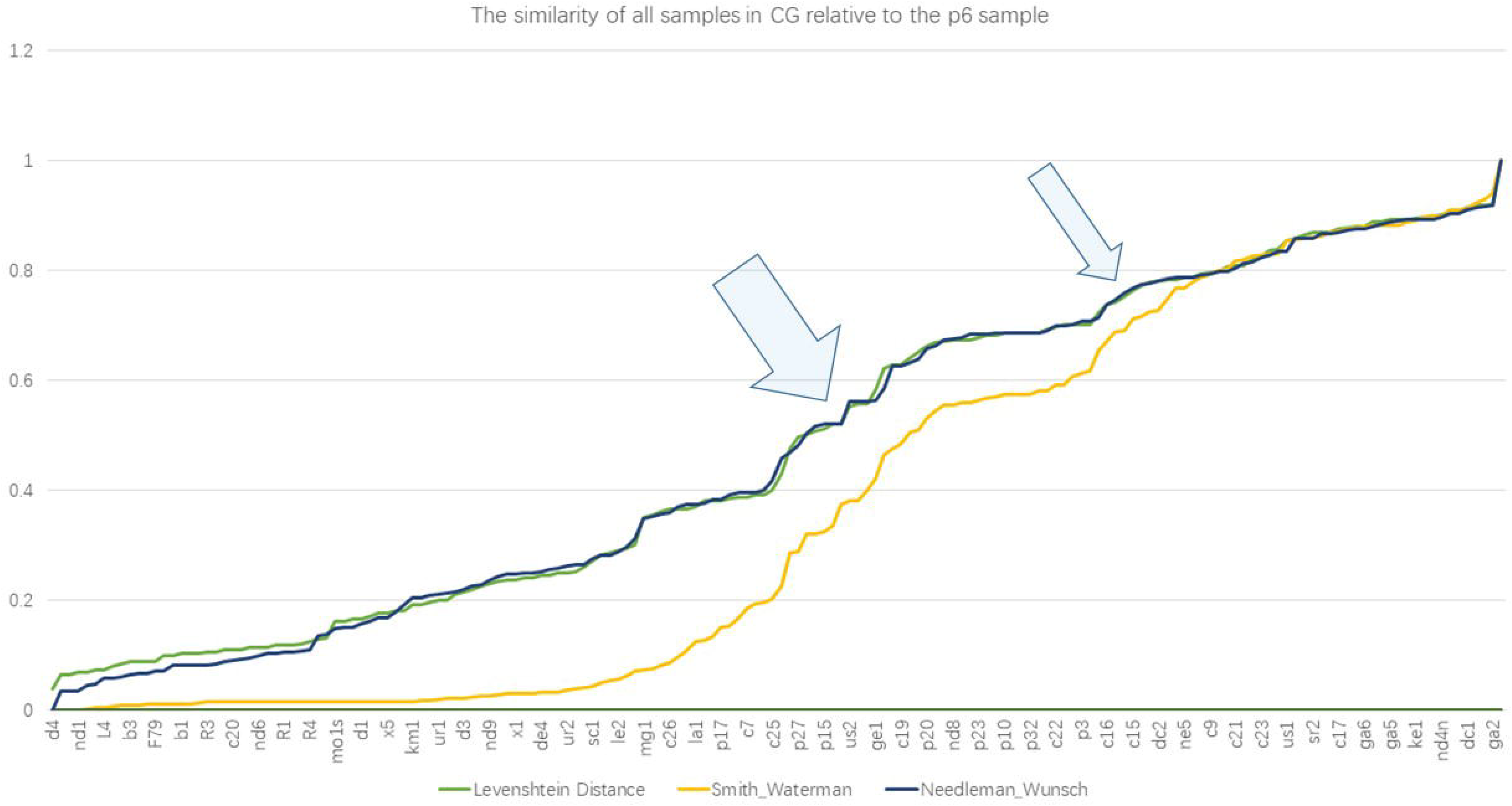
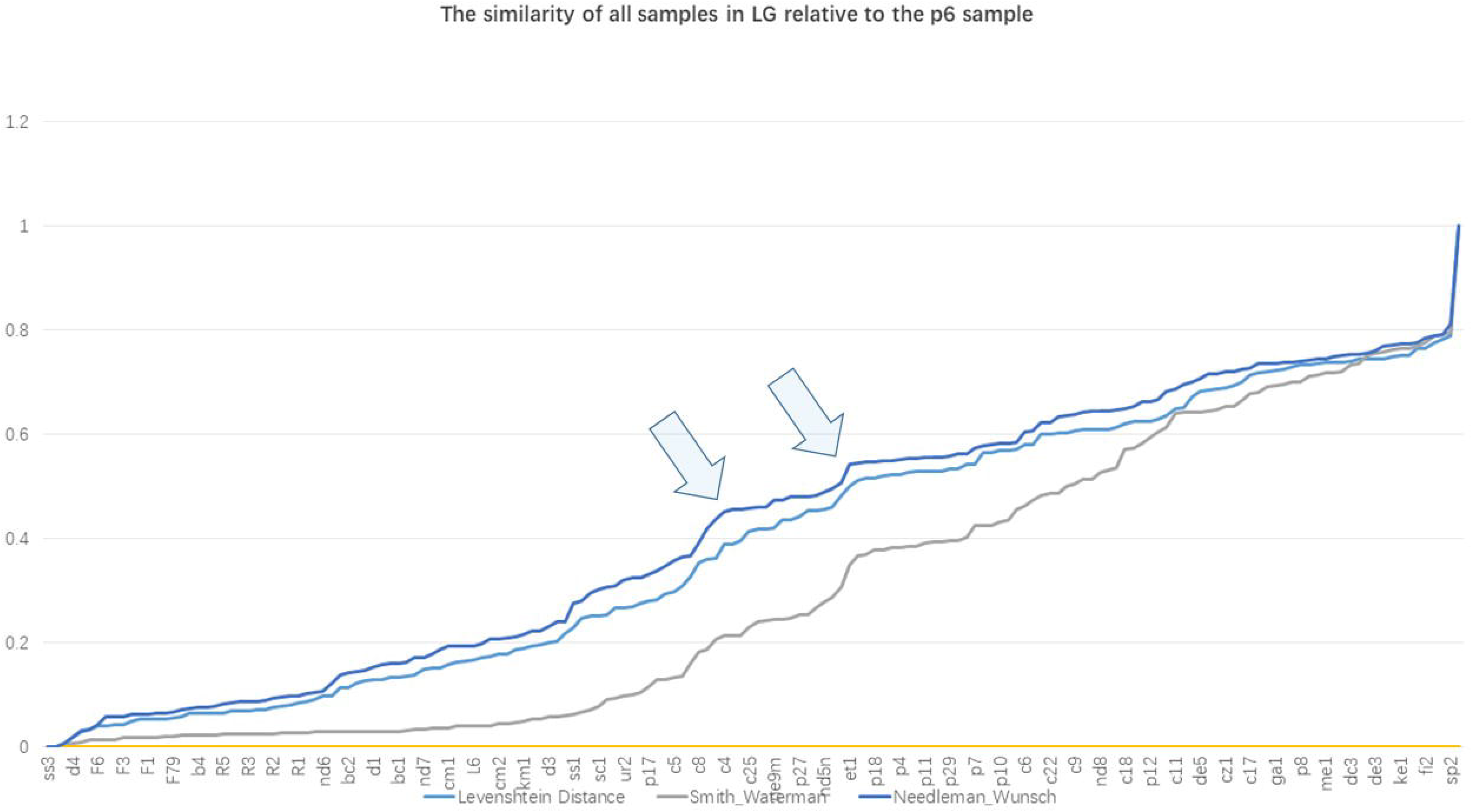
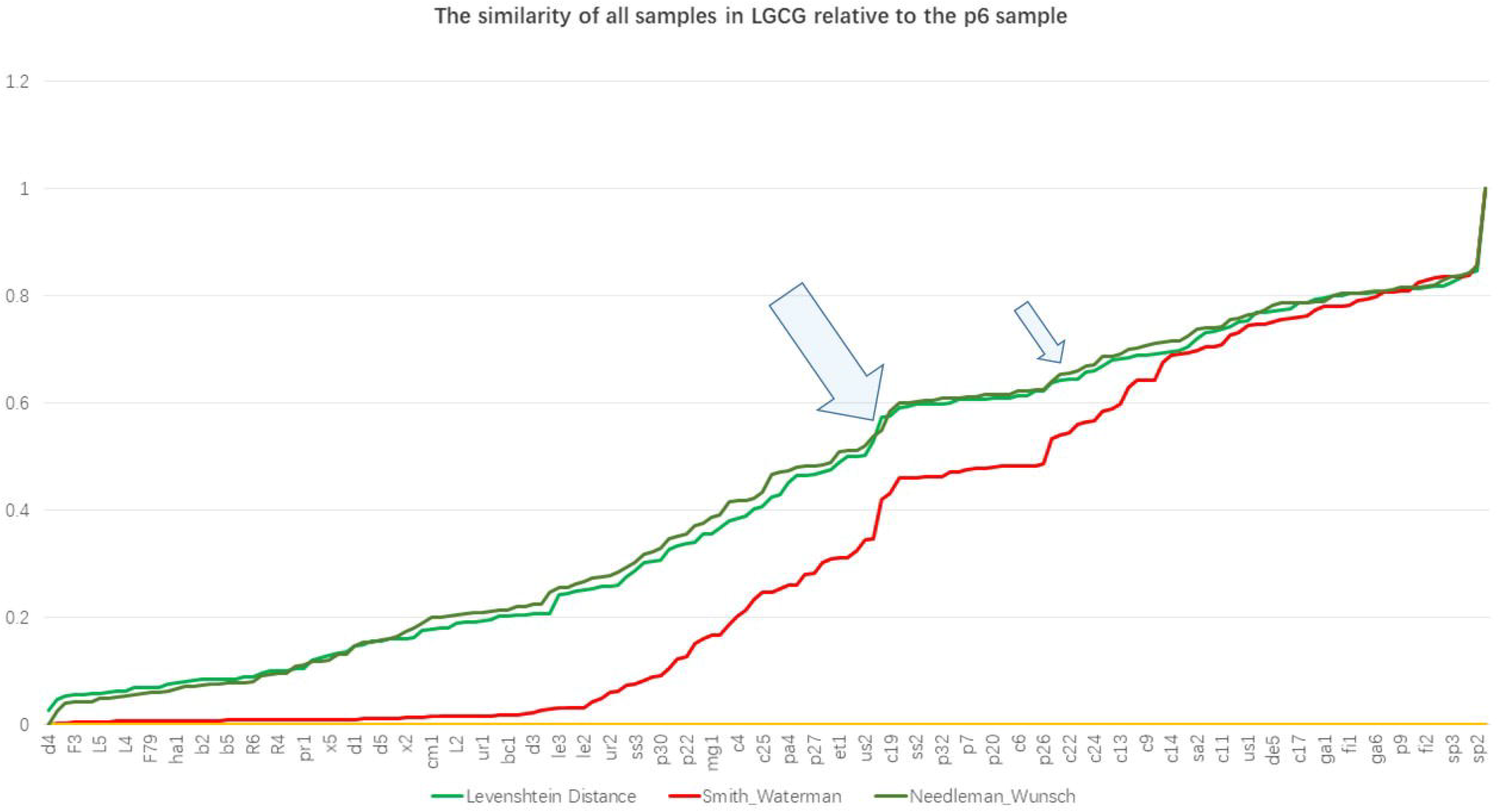
Similarity of SNP profiles among all samples relative to modern human sample p6. (A)CG; (B) LG; (C) CGLG (Detailed sample distribution along the X-axis can be seen in Supplementary file 1∼3)

Hu et al. used a coalescent model to predict past human population sizes from 3154 present-day human genomes [23]. The model detected a reduction in human ancestor population size from about 100,000 to 1280 individuals between around 930,000 and 813,000 years ago. The described bottleneck is congruent with a substantial chronological gap in the available African and Eurasian fossil record, and suggests coincident speciation event. It is interesting to tackle whether such an event was accompanied with the potential LG/CG/LGCG turning points described in this study. Therefore, it will be very valuable to continuously collect fossil DNA information from this period (930,000 and 813,000 years ago) or samples that have a direct genetic connection with this period in the future.

## Conclusion

This study examined the genetic differences of 170 whole-genomes at 239 SNP loci of 18 language genes (LG) and 223 SNP loci of 18 cognitive genes (CG), clustered the SNP data of the above samples through PCA, and calculated the SNP pattern similarity between each sample under three perspectives: LG, CG, and CGLG. The basic conclusions include (1) both the polymorphic patterns of language genes and cognitive genes have undergone different evolutionary stages; (2) the polymorphic patterns of language genes and cognitive genes show significant differences in their early manifestations during human evolution, as reflected in the early patterns of SNP diversity in language genes being very similar across all animal samples, whereas cognitive genes exhibit significant variation right from the start among the selected animal samples; and (3) it seems that samples from all five continents can be seen at every stage of evolution, indicating that new evolutionary populations have always had enough time to spread among the continents.

## Acknowledgments

This study was supported by State Language Commission Research Grant (YB135-117), Association of Chinese Graduate Education Grant (B-2017Y0505-079) and National Research Center for Foreign Language Education Grant (ZGWYJYJJ10A042).

